# Structural studies on *M. tuberculosis* decaprenyl phosphoryl-β-D-ribose epimerase-2 enzyme involved in cell wall biogenesis

**DOI:** 10.1101/2020.10.15.341941

**Authors:** Shanti P. Gangwar, Arkita Bandyopadhyay, Ajay K. Saxena

## Abstract

The *Mycobacterium* DprE2 is a NADH-dependent enzyme and converts the decaprenylphosphoryl-β-D-ribose (DPX) to decaprenylphosphoryl-β-D-arabinofuranose (DPA). The FAD-containing oxidoreductase *MtbDprE1* and NADH-dependent reductase *MtbDprE2* enzymes catalyses together the epimerization reaction, which coverts DPR to DPA. Here, *MtbDprE2* enzyme was purified and structurally characterized using circular dichroism, molecular modelling and dynamics simulation techniques. The *MtbDprE2* was purified, which eluted as oligomer from size exclusion column. The circular dichroism analysis yielded ~ 47.6% α-helix, ~ 19.8% β-sheet and ~ 32.6% random coil structures in *MtbDprE2* enzyme and showed highly thermostability. The molecular modelling of *MtbDprE2* and its complex with NADH showed that it contains two domains (i) the large domain consists of central twisted seven β-sheets decorated by eight α-helices and (ii) a small domain contains two short α-helices connect by loop. Overall, the *MtbDprE2* adopts a typical short-chain dehydrogenase rossmann fold and NADH binds to Asp69, Ser147, Tyr160, Lys164 of catalytic triad and Gly16, Ser19, Glu20, Ile21 of Gly-rich motif of *MtbDprE2*. 1 ns dynamics simulation was performed on apo and NADH bound *MtbDprE2*, which indicated the small conformational change in ligand binding site, which resulted more closed pocket than open pocket observed in apo enzyme. Small conformational changes were observed in active site residues and orientation between large and small domains of *MtbDprE2* upon NADH binding. Current knowledge of *MtbDprE2* structure and its NADH binding mechanism will contribute significantly in development of specific inhibitors against *M. tuberculosis.*

## 1. Introduction

Emergence of multiple drug resistance, extremely drug resistance and total drug resistance strains of *M. tuberculosis* have affected significantly the treatment of tuberculosis **[1]**. In addition, reemergence of infection from dormant stage, slow growth and complex thick cell wall of *M. tuberculosis* has aggravated the tuberculosis infection. The *M. tuberculosis* cell wall contains a unique structural components and lipid layers, which makes it impermeable to many drugs and protects the pathogen from host immune system.

The *M. tuberculosis* DprE1 and DprE2 enzymes are involved in epimerization of DPR to DPA, a precursor required for polysaccharides, arabinogalactan and lipoarabinomannan synthesis in mycobacterial cell wall **[2–3].** The DPA covalently attached the mycolic acid to peptidoglycan and involved in complete cell wall synthesis. The *MtbDprE2* gene knockout study has shown that it is essential for mycobacterial survival **[4]** and requires NADH cofactor for its epimerization reaction **[5].**

The first and second line of tuberculosis drugs inhibit the components of mycobacterial cell wall synthesis **[7].** The BTZ043 and DNB are found highly active against multidrug resistant and extensively drug resistant strains of *M. tuberculosis*. Both series of nitro-aromatic compounds target the heterodimeric decaprenylphosphoryl-β-D-ribose 20-epimerase enzyme system encoded by *MtbDprE1* and *MtbDprE2* genes **[8, 9]**. The BTZ043 has shown low minimal inhibitory concentration ~ 1 ng/ml against *M. tuberculosis*, significantly lower than other tuberculosis drugs **[8].** The BTZ043 covalently modifies the *MtbDprE1*, ablating its function **[10]** and currently undergoing in clinical trial. Other inhibitors of *MtbDprE1* (i) 2-mercaptobenzothiazole **[11]** (ii) 1, 2, 3 triazole conjugates **[12]** (iii) next generation banzothiazinones **[13]** and (iv) non-covalent inhibitors from scaffold morphing [**14]** have been identified and characterized. The crystal structures of several complexes of *Mtb*DprE1 with synthetic inhibitors have been also determined [**15, 16].**

The *Mtb*DprE1 and *MtbDprE2* proteins are essential for mycobacterial growth **[17]** and interacts to each other and considered as two subunit of single enzyme, decaprenyl phosphoryl-β-D-ribofuranose 2-epimerase. In *Corynebacterium glutamicium*, the bacterial two hybrid system has shown the interaction between DprE1 and DprE2 enzymes **[18].** The MtbDprE1 and *MtbDprE2* complex model has been proposed using molecular modeling, threading and dynamics simulation techniques **[19].** The mutational and dynamics simulation analysis on InhA-NADH complex has shown the mechanism of drug resistance in *M. tuberculosis* **[20].**

So far, little is known about structure and mechanism of *MtbDrprE2* enzyme. The analysis of structure and NADH binding of *MtbDprE2* will be critical in development of specific inhibitors against *M. tuberculosis*. In present study, the *MtbDprE2* was purified and secondary structure analysis was performed using circular dichroism spectroscopy. Molecular modeling and dynamics simulation analysis on wild type *MtbDprE2* and its complex with NADH were performed to understand the structural basis of NADH binding to *MtbDprE2*.

## 2. Materials and methods

### 2.1. Expression and purification

*MtbDprE2* gene [*Rv3791*, 254 residues, 27 kD] was cloned in *pET28a* (+) expression vector using NdeI and XhoI restriction sites. Forward (5’- GAT**C**CATAT GATGGTTCTTGATGCCGTA -3’) and reverse (5’- CATGCTCGAGTCAGATGGGCAGCTTGCG- 3’) primers were used for *MtbDprE2* gene amplification from *H37Rv* genomic DNA. The resulting *MtbDprE2* plasmid was checked by restriction-digestion experiment. The *MtbDprE2* plasmid contains 6xHis tag and thrombin cleavage site at N-terminal and 254 residues of *MtbDprE2*. The *MtbDprE2* plasmid was transformation in *Escherichia coli BL21(DE3)* cells, however protein expressed as inclusion body. Different variables, IPTG concentration, temperature, various *E. coli* cell lines were tried, however protein expressed only as inclusion body in cell.

The *MtbDprE2* gene was further cloned in *pET-SUMO* vector (*Invitrogen*) using TA cloning method and checked by gene sequencing. The *E. coli BL21(DE3)* cells were used for *MtbDprE2* plasmid transformation. The 3 l LB media (50μg/ml Kanamycin as antibiotics) was to grow single colony and grown at 37°C, till OD_600_ ~ 0.7-0.8. 0.25 mM IPTG was used to induce the culture at 37°C and grew further for 3 hr. The culture was centrifuged at 10,000 x g, collected the pellet and suspended in buffer-A (20 mM Tris/HCl pH 8.0, 5% (v/v) Glycerol, 300 mM NaCl, 1 mM Benzamidine-HCl, 1 mM Phenylmethylsulfonyl fluoride, 2 mM β-mercaptoethanol and 0.3 mg/ml Lysozyme). The culture was kept on ice for 1 h, sonicated and supernatant was collected by centrifugation at 18,000 x g for 20 min.

The Ni-NTA column was pre-equilibrated with buffer-B (20 mM Tris/HCl pH 8.0, 300 mM NaCl, 5% Glycerol, 1 mM Phenylmethylsulfonyl fluoride, 1 mM Benzamidine-HCl and 2 mM β-mercaptoethanol) and loaded the supernatant. The column was washed with buffer-B +15mM Imidazole and eluted the protein in buffer-B+ 300mM imidazole. The peak fractions were pooled, concentrated and loaded on Superdex 75(16/60) column (*GE Healthcare*) preequilibrated in buffer 20 mM Tris/HCl pH 8.0, 300 mM NaCl and 5% Glycerol. We pooled the peak fractions and used ultracentrifugal device (*Amicon*) to concentrate the protein.

The SUMO protease (a ubiquitin like protein processing enzyme) was used to cleave SUMO tag from *MtbDprE2*-SUMO fusion protein. 200 μg of *MtbDprE2*-SUMO fusion protein was dissolved in 500 μl of SUMO protease in buffer (500mM Tris/HCl pH 8.0, 1.5 M NaCl, 2% NP-40, 10mM DTT) and 2 μg of SUMO protease was added in it. The reaction mixture was incubated at 4 °C and aliquots were taken out after 4, 8, 16 and 24 h. The aliquots were examined on SDS-PAGE to check the cleavage reaction. The cleaved *MtbDprE2* was further purified using Superdex 75(16/60) column. The purity of MtbDprE2 was analyzed on SDS-PAGE and estimated the concentration using Bradford method.

### 2.2. Circular dichorism analysis

Chirascan™ CD spectropolarimeter (*Applied Photophysics*) was used to collect the CD data on *MtbDprE2* in 260-200 nm wavelength. The *MtbDprE2* was concentrated to 0.1 mg/ml in b10mM Tris/HCl buffer pH 8.0 buffer and loaded on 1-mm sample cuvette for CD data collection. The protein buffer was used as blank and subtracted from each reading. Three sequential scans were collected for each data. The mean residue ellipticity (deg.cm^2^/dmol) was calculate using Dichroweb server **[21].** For thermal denaturation analysis, the CD data on *MtbDprE2* was collected from 20°C to 70°C in 10°C interval. Before measurement, the sample cuvette was incubated at each temperature and checked with water bath.

Various theoretical structural prediction programs e. g. Jpred **[22],** Raptor X **[23],** HNN **[24],** DSC **[25],** PHD **[26],** GOR **[27],** CFSSP **[28]** and SOPMA **[29]** were used to calculate the secondary structure of *MtbDprE2*. The PSIPRED **[30]** analysis on *MtbDprE2* sequence is shown in Fig. 2A.

### 2.3. Homology modeling

The ITASSER server **[31]** was used to obtain the *MtbDprE2* model (1-254 residues). LOMETS software **[32]** used the *MtbDprE2* sequence and identified the best template from protein database. The SPICKER program **[33]** identified the best template with high Z-score (high threading alignment score) after simulation of structure assembly. Based on the C-score, the program predicts the best score. The TM-align program yielded the top 10 models having best TM-score. PROCHECK **[34]** program was used analyzed the quality of *MtbDprE2* model.

### 2.4. MtbDprE2-NADH docking analysis

The PDB-4JRO [Crystal structure of FabG+NADP+ complex from *Listeria monocytogenes*] was used as template to dock NADH into *MtbDprE2* model. To optimize NADH docking, the GLIDE program **[35]** of Schrödinger-9.4 suite was used with IFD (induced fit docking) module. We used the XP (Extra Precision) scoring function and scaled the van der Waals radii of *MtbHddA* by 0.6 fold. For prime site optimization, all *MtbDprE2* residues located within 4 Å radii of NADH were refined. The Glidestone module of Glide program was used to obtain the best *MtbDprE2*+NADH complex using 5,000 cycles of scoring and 5,000 cycles of minimization. The *MtbDprE2*+NADH complex having lowest IFD value was selected as the best *MtbDprE*2+NADH *c*omplex.

### 2.5. Dynamics simulation analysis

The dynamics simulation on apo-*MtbDprE2* and its complex with NADH were performed using GROMACS 2020.1-MODIFIED version **[36]** taking OPLSAA force field **[37].** The protein was kept in cubic box having 0.6 nm spacing and filled with TIP3P water molecules. The entire charge of system was neutralized by adding chloride and sodium atoms and periodic boundary condition was used for dynamics simulation. The system was subjected to 5000 steps of steepest descent minimization. Before dynamics, the solvent and ions were equilibrated surrounding the protein. Initially, first phase of NVT equilibration (100 ps) was performed, which stabilize the temperature of the system. In second phase, equilibration of pressure was conducted in 100 ps NPT equilibration (constant number of particles, pressure and temperature).

A constant temperature 300 K and coupling constant (τ) ~ 0.1 ps was maintained by applying V-rescale (modified Berendsen thermostat) coupling with coulomb cutoff 1. The 1 atm pressure was maintained, isotropic scaling and 2 ps relaxation time. LINCS algorithm was used to constrain the bond lengths. The long ranger electrostatic interactions was calculated using PME method **[38]**. A force constant of 1000 kJ/mol.nm^2^ was used to restrain the protein harmonically. 1 ns molecular dynamics simulation with time step ~ 2 fs was performed and saved the coordinates after every 5 ps for trajectory analysis. The least square fitting procedure was used to calculate RMSD and RMSF and plotted by Plot2X program **[39].** PROCHECK program **[34]** was used to check the stereochemistry of simulated *MtbDprE2* models and Chimera program **[40]** for structure visualization.

## 3. Results and discussion

### 3.1. Purified MtbDprE2 exists as oligomer

*MtbDprE2* sequence (254 residues, 27 kDa) is shown in Fig. 1A and residues involved in NADH binding are shown in blue letters. The *MtbDprE2* gene was cloned into *pET28*(a+) expression vector, however protein expressed as inclusion body in *E. coli. BL21(DE3)* cells. Various parameters e. g. IPTG concentrations (0.1-1mM), different temperature (10 - 37 °C) and different cells (*E. coli* arctic, *BL21-codon-Plus*) were tried, however protein expressed insoluble fraction of cell. The *MtbDprE2* gene was further cloned in *pET-SUMO* vector using TA cloning method and clones were confirmed restriction-digestion (Fig. 1B). The *E. coli*. *BL21(DE3)* cells were used for *MtbDprE2* plasmid transformation and protein overexpressed as soluble protein. The *MtbDprE2* fusion protein eluted as oligomer from Superdex 200(16/60) column. The *MtbDprE2* fusion protein was treated with SUMO protease and examined on SDS-PAGE (Fig. 1C, inset). The SDS-PAGE analysis showed the fusion protein, cleaved DprE2 and cleaved SUMO tag (Fig. 1C, inset). The cleaved *MtbDprE2* was further purified on Superdex 200(16/60) column, which eluted as oligomer from column (Fig. 1C).

**Fig. 1.**
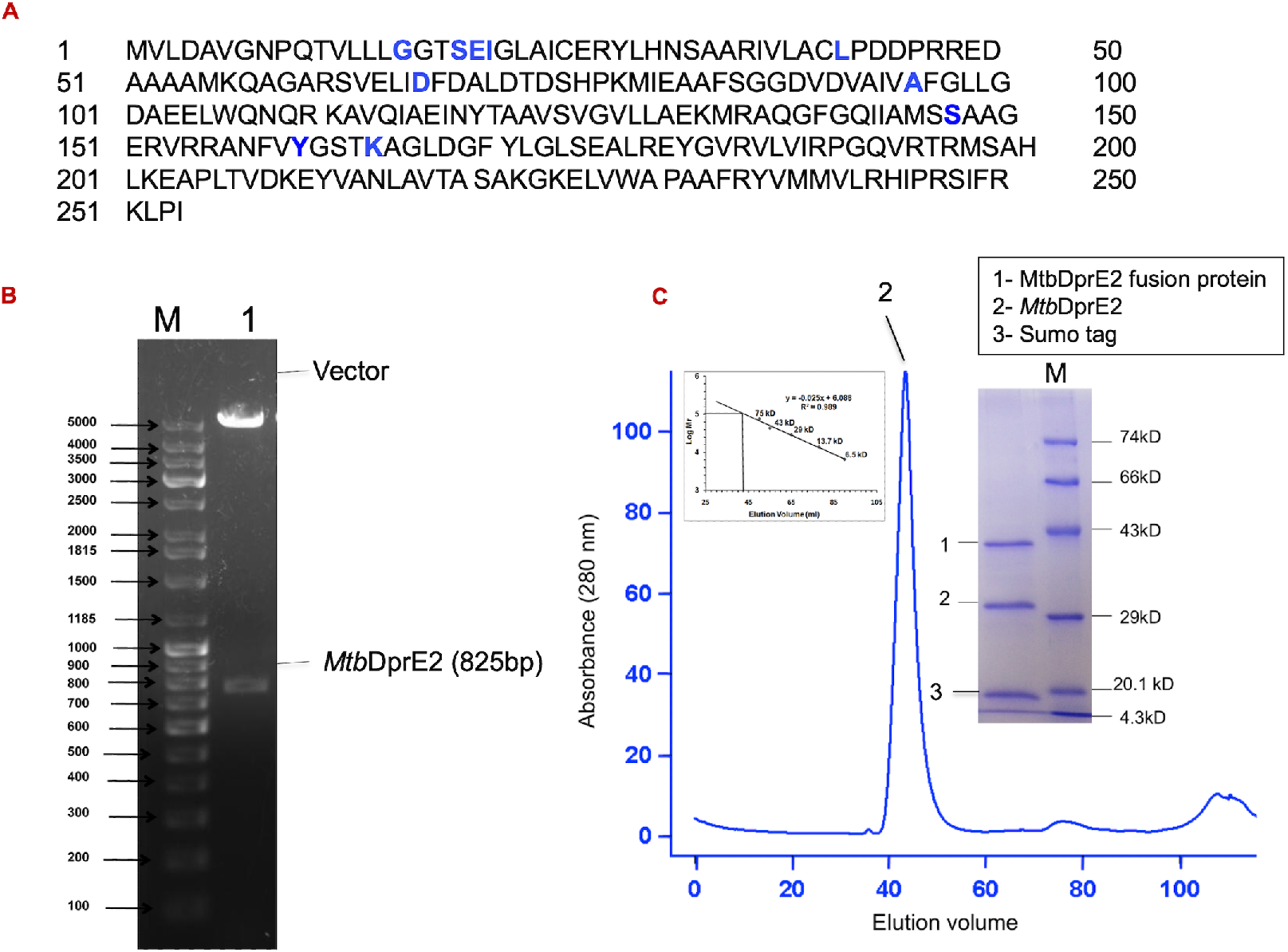
**A,** *MtbDprE2* sequence showing the residues involved in NADH binding (blue) and in substrate-binding (red). **B,** Restriction-digestion analysis of SUMO-*MtbDprE2* plasmid showing the *MtbDprE2* gene fall out. **C,** SDS-PAGE analysis of SUMO-*MtbDprE2* fusion protein after cleavage with SUMO protease (1- SUMO and 2- *MtbDprE2*). The cleaved *MtbDprE2* eluted as oligomer from Superdex 200(16/60) column, as identified using molecular mass standard.

### 3.2. Secondary structure and thermal denaturation analysis

The PSIPred analysis **[30]** on *MtbDprE2* sequence (Fig. 2A) showed seven β-strands and ten α-helices structures in protein. The CD data was collected in 260-200 nm wavelength range and secondary structure was calculated using DICHROWEB server **[41].** The program yielded ~ 47.6% α-helix, ~ 19.8% β-sheet and ~ 32.1% random coil in *MtbDprE2* structure (Fig. 2B). Theoretical structure prediction on *MtbDprE2* also yielded quite similar secondary structure, as observed in CD data (Table 1). For thermal denaturation analysis, CD data on *MtbDprE2* was collected from 20 °C −70 °C in 10 °C step (Fig. 2C). These data showed that minor changes in secondary structures, indicating high thermostability of *MtbDprE2* enzyme.

**Figure 2.**
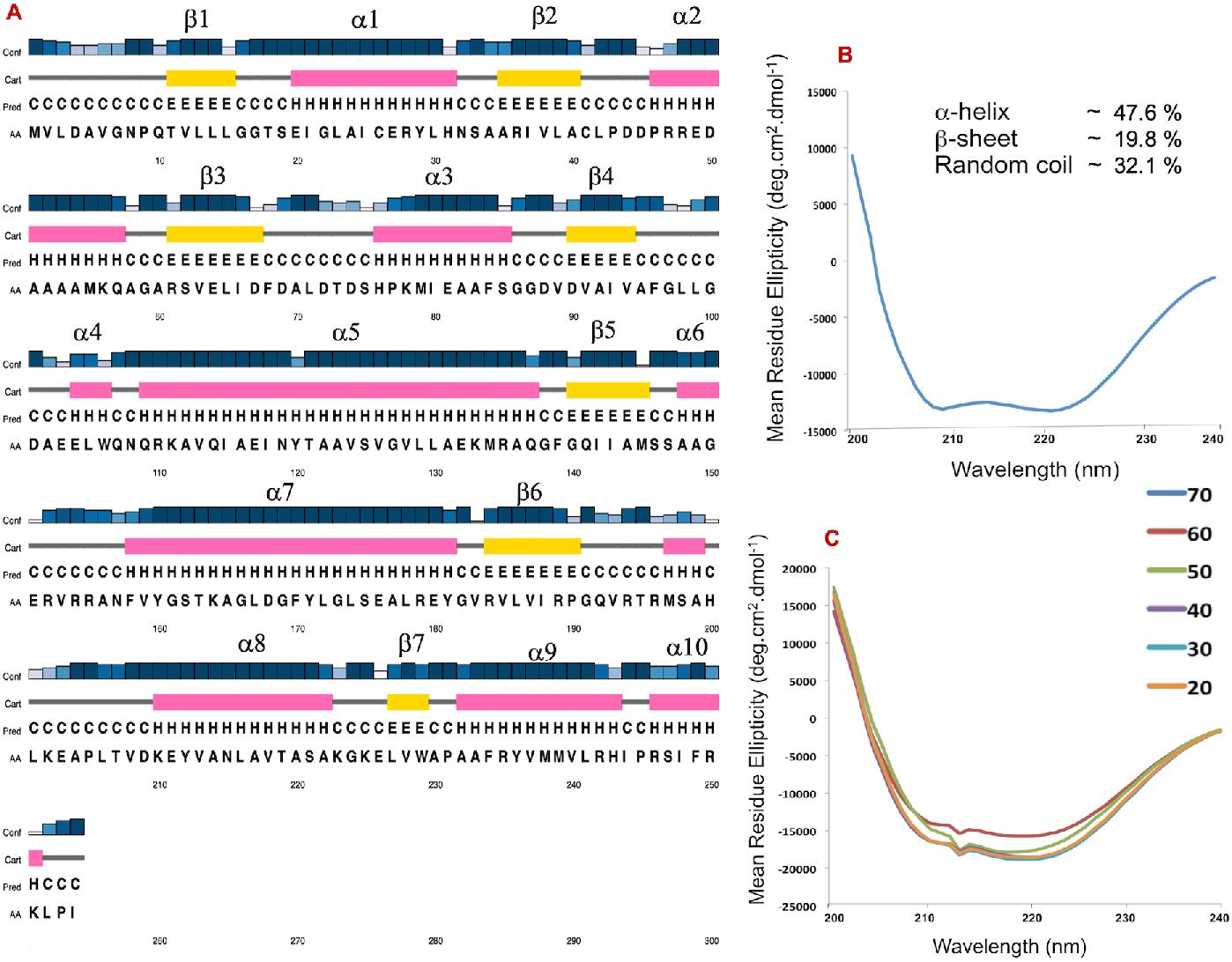
Secondary structure and thermal denaturation analysis of *MtbDprE2* using CD spectroscopy. **(A)** The PSIPRED analysis showing the secondary structural contents of *MtbDprE2*. The α-helices are shown magenta spiral and β-strands in yellow arrow. **(B)** The 260 to 200 nm wavelength was used for CD data collection on *MtbDprE2* and secondary structures were estimated by Dichroweb server **[41].** *Inset* shows the *MtbDprE2* secondary structure **(C)** The CD data collected 260-200 nm range starting from 20 °C to 70 °C range with 10 °C interval.

**Table 1.** Secondary structural composition of *MtbDprE2* obtained from circular dichroism and theoretical structure prediction programs

| **Programs** | **α-helix** | **β-sheet** | **Random coil** |
| --- | --- | --- | --- |
| **CD** | **47.6%** | **19.8%** | **32.6%** |
| SPOMA18 | 46.5% | 25.2% | 28.3% |
| GOR4 | 54.3% | 11.4% | 34.2% |
| PHD | 46.5% | 16.1% | 37.4% |
| DSC | 40.5% | 15.7% | 43.7% |
| HNN | 53.5% | 15.7% | 30.7% |
| RAPTOR | 44% | 12% | 42% |

### 3.3. Molecular modeling

The I-TASSER server was used to obtain the *MtbDprE2* model (1-254 residues). The server identified the PDB-4JRO (Crystal structure of FabG + NADP+ complex from *Listeria monocytogenes*) as the best template with following parameters *e.g.* (id1= 0.20, id2 = 0.24, cov = 0.93, Z-score = 2.07). The best *MtbDprE2* model was obtained with following parameters [C-score = −0.74 and TM-score = 0.62± 0.14]. The PROCHECK program showed the good stereochemistry of obtained *MtbDprE2* model and all residues lie in allowed ϕ, ψ regions of Ramachandran plot (**Fig. S1**). The ProSA **[42]** and ERRAT **[43]** servers analysis on *MtbDprE2* model also showed the good stereochemistry and high reliability **(Fig. S1).**

The *Mtb*DprE2 model belongs to short chain dehydrogenase/ reductase (SDR) superfamily of enzymes with Rossman fold containing NADH/NAD(P)H binding site. The *MtbDprE2* model contains two domains, (i) the major domain, which adopts a typical rossman-fold with seven stranded β-sheets surrounded by eight-α helices and (ii) a small domain contains two α-helices (a6-a7) connected by a loop (**Fig. 3A**). The electrostatic surface diagram of *MtbDprE2* contoured at ±10 kT (**Fig. 3B)** showed that NADH binding site contains partially positively charge, while overall surface of enzyme is more negatively charge and partial positive charges are scattered all over the surface.

**Figure 3.**
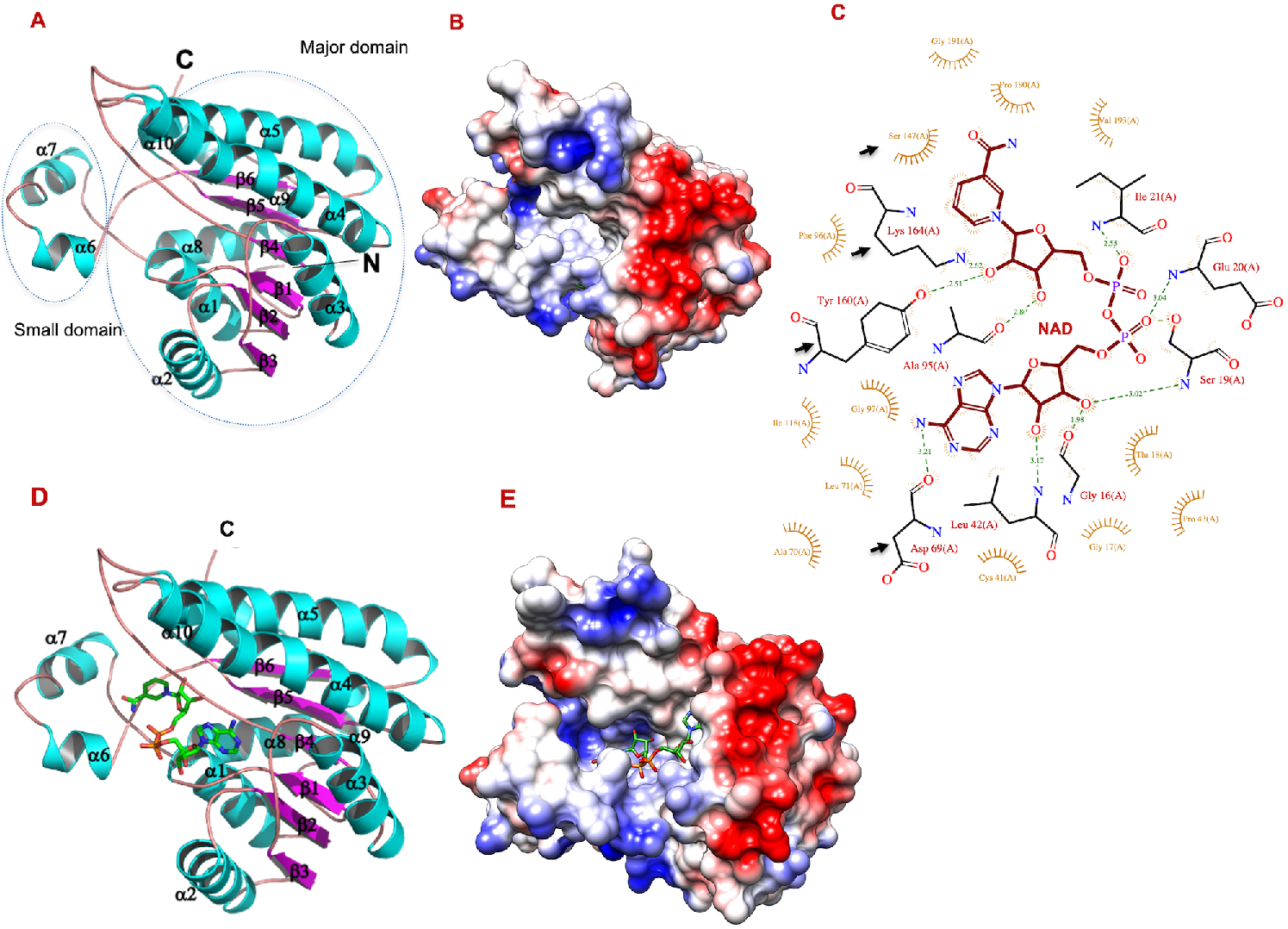
**(A)** The *MtbDprE2* model (1-254 residues) obtained from I-TASSER server **[37].** The α–helices in cyan, β–sheets in magenta and loops are shown in light orange colors respectively. **(B)** Electrostatic surface diagram of *MtbDprE2* contoured at −10kT to +10kT. Overall surface of *MtbDprE2* is negatively charged, while positive charges spread around the surface. The NADH cofactor binding site is partially positively charged. **(C)** The LigPlot **[52]** showing the interactions between NADH and *MtbDprE2*. Hydrogen bonds are shown in green dashed lines and van der waals interactions in orange Arc color **(D)** Stereogram of *MtbDprE2*+NADH complex model. The NADH is shown in green color. (E) Electrostatic surface diagram of *MtbDprE2* contoured at −10kT to +10kT with NADH fitted in the binding pocket.

### 3.4. NADH binding analysis

The PDB-4JRO was used as template to dock NADH into *MtbDprE2*. Obtained *MtbDprE2*+NADH complex was used as template in GLIDE program of schrÖdinger suite to optimize the fitting of NADH into *MtbDprE2* and induced fit docking protocol was used for it.

The LigPlot **[44]** analysis showed that NADH occupies the deep cleft of *MtbDprE2* and forms hydrogen bonds with Asp69, Ser147, Tyr160, Lys164 residues of catalytic triad, Gly16, Ser19, Glu20, Ile21 residues of Gly-rich motif and Leu42, Ala95 residues of *MtbDprE2* (Fig. 3C). In SDR family members, the active site is usually formed by Tyr-X-X-X-Lys sequence **[45, 46].** In *MtbDprE2*, Tyr160-Gly161-Ser162-Thr163-Lys164 motif is found in catalytic site, as observed in ‘classical’ type of SDR superfamily. The Asn99 residue of *MtbDprE2* is highly conserved to SDR family of proteins, however, does not interact to NAD in *MtbDprE2*. Instead, highly conserved Asp69 interacts with NAD and forms the catalytic triad. Lys164 forms hydrogen bond with NAD nicotinamide ribose and decreased the p*K*a of Tyr160-OH. The Tyr160 catalyzes and Ser147 immobilizes the substrate and thus forms the Ser147, Tyr160, Lys164 catalytic triad.

The TG-xxxGxG consensus sequence in SDR family enzymes is involved in NADH binding **[47].** In *MtbDprE2*, the Gly rich motif L15-**G16**-G17-T18-**S19-E20-I21**-G22-L23-A24 (shown in #) is observed at first β1-strand. The Gly16 residue of *MtbDprE2* is highly conserved, while S19, E20 and I21 residues are partially conserved and interacts with 2′-phosphate and ribose hydroxyl groups of NADH.

The β4-α4 loop of *MtbDprE2* contains ^91^NNAGX^95^ motif, which stabilizes the central α6 helix and β-sheet, which contains two catalytic residues (Tyr157 and Lys161) **[48, 49].** In *MtbDprE2*, ^90^DVAIVAFGL^99^ motif represents the same loop though conserved, but may play a role like ^91^NNAGX^95^ motif, as found in other short chain dehydrogenase enzyme.

In apo-*Mtb*DrpE2, the NADH binding pocket is quite wide (Fig. 3B), compared to ligand bound, as surrounding helices tilted outwards from the pocket binding (Fig. 3E). The *MtbDprE2* showed little conservation and similarity with other members of SDR family of enzymes and pocket involved in substrate binding has shown high degree of variability **[50].**

### 3.5. Sequence alignment and comparative structure analysis

The *MtbDprE2* structure was aligned with structures of PDB database using TM-structural alignment program of I-TASSER server, which yielded ten closest structural homologs (as shown in Table 2), (i) PDB-3awd, identity=17.4%, 97.2% coverage **[51]** (ii) PDB-3s55, identity=18.4%, 96.5% coverage **[52]** (iii) PDB-1gee, identity=15.4%, 96.9% coverage (iv) PDB-2uvd, identity =15.6%, 95.7% coverage **[53]** (v) PDB-3aut, identity=16.6%, coverage=97.2% **[54]** (vi) PDB-3t7c, identity=13.3%, coverage=97.2% **[55]** (vii) PDB-1ipe, identity=18.1%, coverage=97.6% **[56]** (viii) PDB-2wsb, identity=18.4%, coverage=96.5% **[57]** (ix) PDB=2uve, identity=15 %, coverage=97.2% **[58]** (x) PDB-4nbv, identity=21%, Coverage=95.7% **[59].**

**Table 2.** Top 10 best structural analogs of *MtbDprE2* from PDB

| Rank | PDB hits | TM-score<sup>a</sup> | RMSD<sup>b</sup> | IDEN<sup>c</sup> | Cov<sup>d</sup> |
| --- | --- | --- | --- | --- | --- |
| 1 | 3awdA | 0.874 | 2.62 | 0.174 | 0.972 |
| 2 | 3s55A | 0.874 | 2.47 | 0.184 | 0.965 |
| 3 | 1geeA | 0.873 | 2.49 | 0.154 | 0.969 |
| 4 | 2uvdH | 0.873 | 2.54 | 0.156 | 0.957 |
| 5 | 3autA | 0.872 | 2.60 | 0.166 | 0.972 |
| 6 | 3t7cA | 0.871 | 2.58 | 0.133 | 0.972 |
| 7 | 1ipeA | 0.870 | 2.58 | 0.181 | 0.976 |
| 8 | 2wsbA | 0.870 | 2.57 | 0.184 | 0.965 |
| 9 | 3uveA | 0.869 | 2.62 | 0.150 | 0.972 |
| 10 | 4nbvA | 0.869 | 2.51 | 0.210 | 0.957 |

(a) TM-score- Ranking of protein
(b) RMSD- RMSD between structural analogs and *MtbDprE2* residues
(c) IDEN- sequence identity
(d) Cov- coverage of the alignment by TM-align

Sequence alignment of *MtbDprE2* with ten structural homologs (Fig. 4A) showed that Asp69, Ser147, Tyr160, Lys164 of catalytic triad (shown as *) are highly conserved in all ten sequences. In addition, Gly16, Ser19, Glu20, Ile21 of Gly-rich motif of *MtbDprE2* are also fully and partially conserved. Two additional residues, Leu44, Ala95 (shown as ^+^) are least conserved in ten sequences. Overall, residues of NADH binding pocket of *MtbDprE2* were quite conserved in all ten homologs, despite having low sequence identity (13-21%) (Table 3). Superposition of structures of ten homologs on *MtbDprE2* has yielded RMSD ~ 0.91 for 184 Ca atoms, indicating quite conserved structures in all sequences (Fig. 4B). Major differences were observed at N- and C-terminal regions, small domain comprising α6–α7 helices and in a small loop (shown in circle) in *MtbDprE2* and ten homologs, while remaining structure was quite conserved.

**Table 3.**
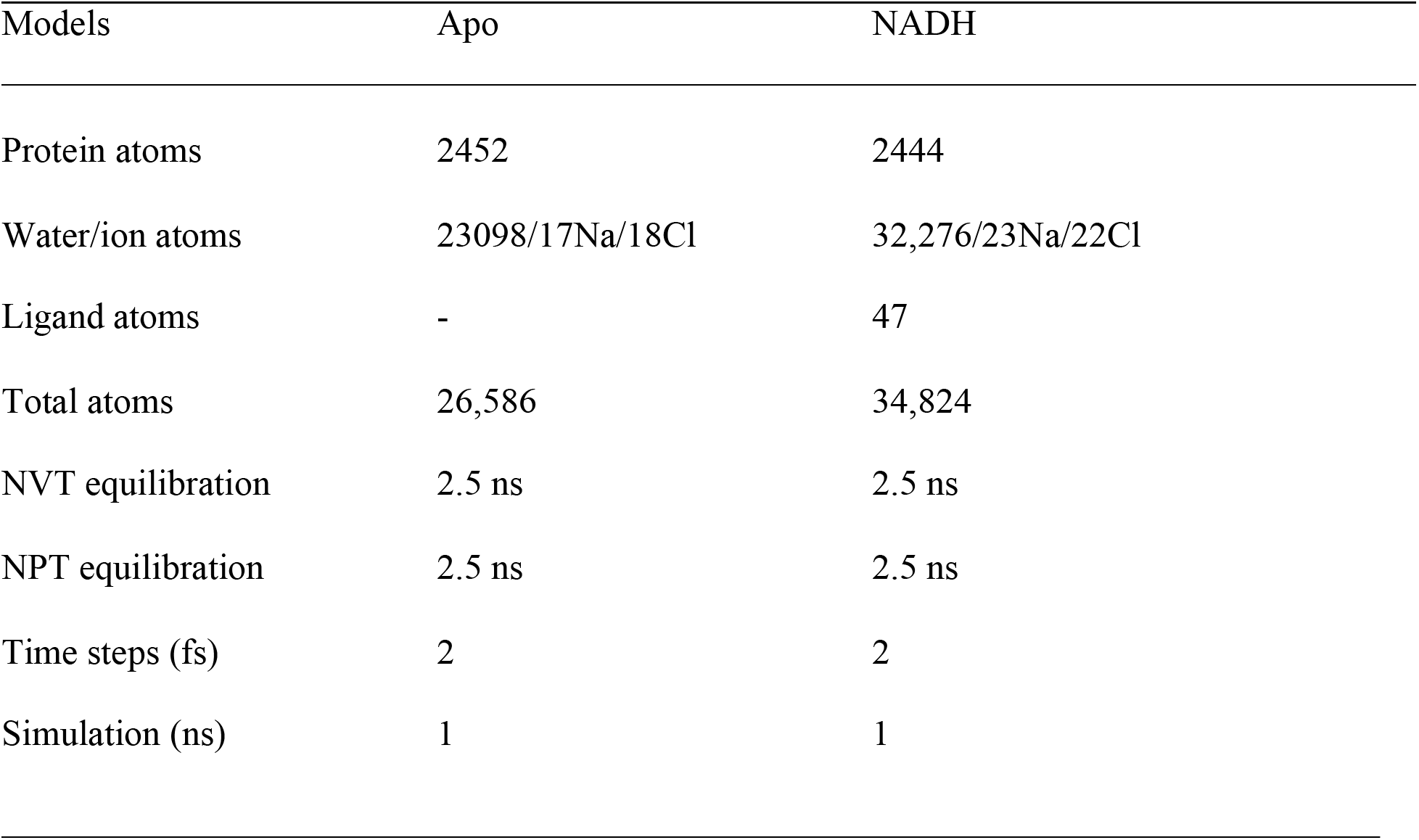
Dynamics simulation on apo-*MtbDrpE2* and its complex with NADH

| Models | Apo | NADH |
| --- | --- | --- |
| Protein atoms | 2452 | 2444 |
| Water/ion atoms | 23098/17Na/18Cl | 32,276/23Na/22Cl |
| Ligand atoms | - | 47 |
| Total atoms | 26,586 | 34,824 |
| NVT equilibration | 2.5 ns | 2.5 ns |
| NPT equilibration | 2.5 ns | 2.5 ns |
| Time steps (fs) | 2 | 2 |
| Simulation (ns) | 1 | 1 |

**Figure 4.**
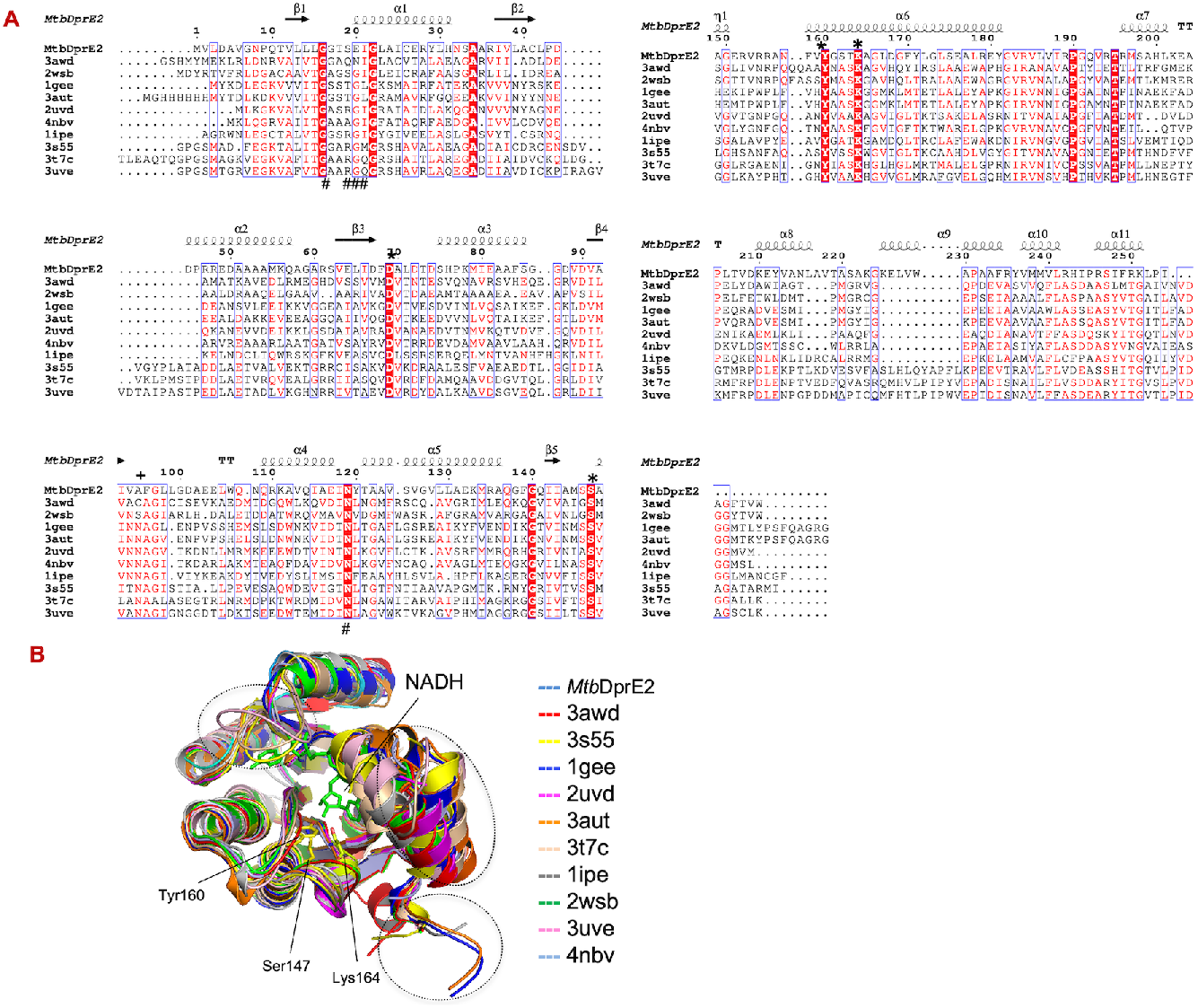
**(A)** Sequence alignment of *MtbDprE2* sequence with ten closest structural homologs obtained from PDB database, (i) PDB-3awd (red) (ii) PDB-3s55 (yellow) (iii) PDB-1gee (blue) (iv) PDB-2uvd (magenta) (v) PDB-3aut (orange) (vi) PDB-3t7c (wheat) (vii) PDB-1ipe (grey) (viii) PDB-2wsb (green) (ix) PDB=2uve (light pink) (x) PDB-4nbv (light blue) using MultiAln and ESPript programs. The secondary structures of *MtbDprE2* model are shown above sequence alignment. The highly conserved residues are shown in red shade and semi conserved in red letters. The NADH binding residues are indicated in (*) above sequence alignment. (**B)** Superposition of (i) PDB-3awd (red) (ii) PDB-3s55 (yellow) (iii) PDB-1gee (blue) (iv) PDB-2uvd (magenta) (v) PDB-3aut (orange) (vi) PDB-3t7c (wheat) (vii) PDB-1ipe (grey) (viii) PDB-2wsb (green) (ix) PDB=2uve (light pink) (x) PDB-4nbv (light blue) on *MtbDprE2* structure (cyan) using PyMol program **[22].** The small domain of *MtbDprE2* having α6 and α7 helices showed large conformational changes (shown in circle) and minor conformational changes were observed in major domain. Residues involved in NADH binding are shown in stick diagram.

### 3.5. Dynamics simulation on wild type MtbDprE2 and its complex with NADH

To examine the *MtbDprE2* enzyme dynamics involved in NADH binding, we performed dynamics simulation on apo and NADH bound *MtbDprE2* and analyzed the trajectories obtained during simulation.

#### Dynamics simulation on apo-MtbDprE2

1 ns dynamics simulation was performed on apo-*MtbDprE2* and analyzed the conformational changes occurred during simulation. The simulated *Mtb*DrprE2 structure (red) was superposed on starting structure (grey), which yielded RMSD ~ 2.5 Å for 240 Cα atoms (Fig. 5A). Overall *MtbDprE2* structure superposed well, except minor changes were observed in loop regions and helical orientation. Major changes in RMSD occurred during ns simulation and remained stable to ~ 0.25 nm during 1 ns simulation (Fig. 5D). Radius of gyration, Rg ~ 1.82±0.20 nm was observed during 1 ns simulation (Fig. 5E). To examine, the structural changes in *MtbDprE2*, we computed the RMSF of *MtbDprE2* residues (Fig. 5F). High RMSF were obtained for N- and C-terminal regions, while remaining structure showed RMSF between 0.1-0.2 nm. The Asp69, Ser147, Tyr160, Lys164 of catalytic triad and Gly16, Ser19, Glu20, Ile21 of Gly-rich motif, involved in NADH binding were quite stable during dynamics simulation.

**Fig. 5.**
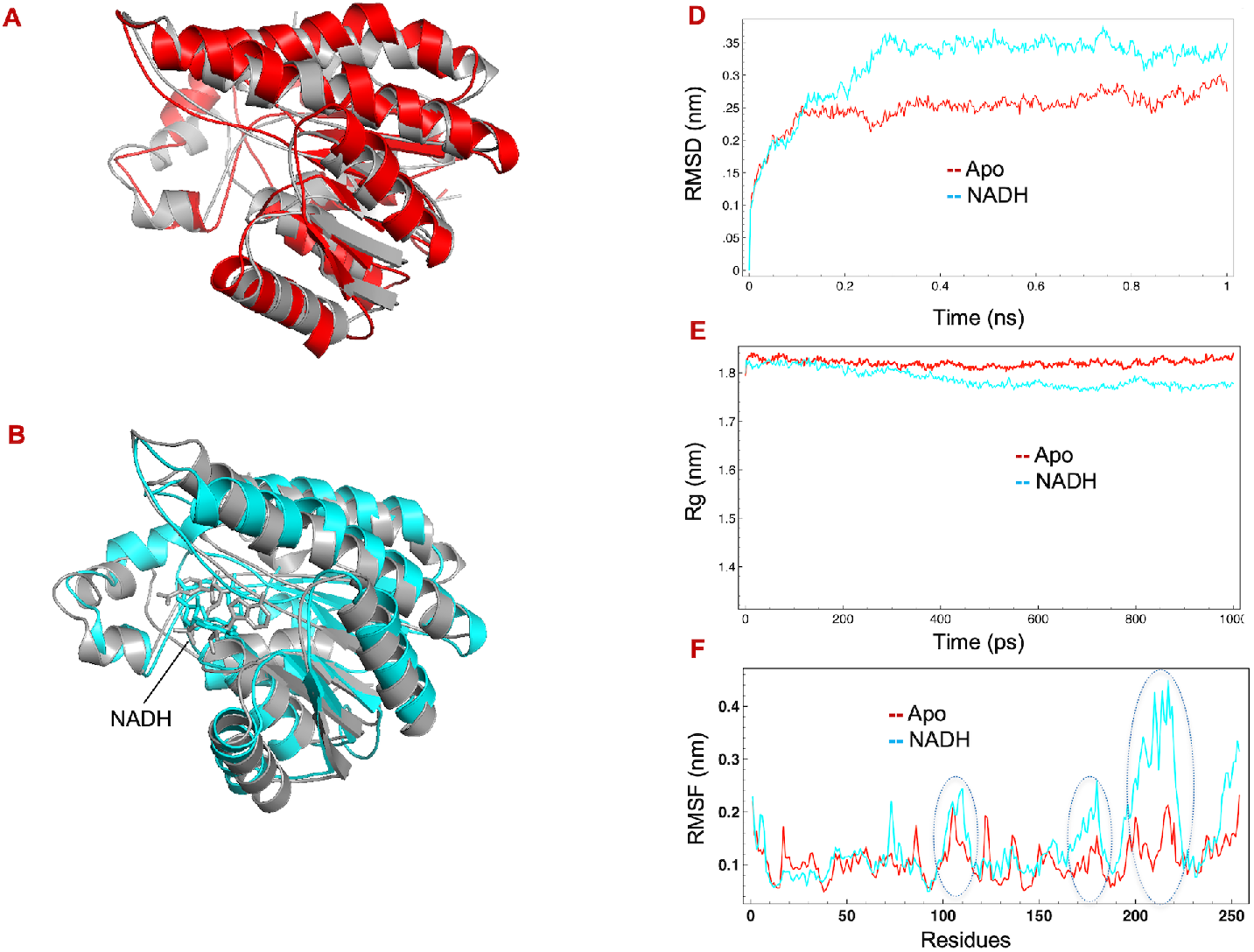
Molecular dynamics simulation on *MtbDprE2* and its complex with NADH **(Table 3). A,** Simulated *MtbDprE2* model (red) is superposed initial *MtbDprE2* model (grey). **B,** Simulated *MtbDprE2*+NADH model (cyan) is superposed on initial *MtbDprE2*+NADH model (grey). **C,** Conformational changes in NADH binding residues after dynamics simulation, initial (red) and simulated (cyan). **D,** RMSD of the backbone Ca atoms (in nm) obtained 1 ns simulation of apo and complexed *MtbDprE2*. **E,** Radius of gyration (in nm) of apo and complexed *MtbDprE2* during 1 ns simulation. **F,** The root square fluctuation (RMSF in Å) of backbone Cα atoms of apo and complexed *MtbDprE2* during 1 ns simulation.

#### Dynamics simulation with MtbDprE2+NADH complex

1 ns dynamics simulation was performed on *MtbDprE2*+NADH complex to analyze the enzyme dynamics involved in NADH binding. The simulated complex structure (cyan) was superposed on starting structure (grey), which yielded RMSD ~ 2.49 Å for 224 Cα atoms (Fig. 5B). The NADH binding inducing small conformational change in NADH binding pocket, which result in more closed pocket than open pocket observed in wild type *MtbDprE2*. The orientation between large and small domains of *MtbDprE2* domains also changes slightly after dynamics simulation (Fig. 5B).

Major changes in RMSD occurred during 0.2 ns simulation and then remained stable to ~ 0.35 nm throughout 1 ns simulation. The radius of gyration, Rg ~ 1.78±0.3 nm was observed during entire simulation (Fig. 5E). To examine the structural changes, we computed the RMSF for each residue of *MtbDprE2* (Fig. 5F). High RMSF values were observed for N- and C-terminal regions, 110-120 residues, 180-190 residues and 200-225 residues of *MtbDprE2*. Residues involved in NADH binding did show high RMSF values in dynamics simulation of *MtbDprE2*+NADH complex.

## 4. Conclusion

The *MtbDprE2* enzyme is involved in DPA biosynthetic pathway and critical for *M. tuberculosis* drug development. Here, we have purified and structurally characterized the NADH binding mechanism of *MtbDprE2*. The CD analysis showed the secondary structures of *MtbDprE2*, quite similar to SDR family of enzymes. The thermal denaturation profile of *MtbDprE2* indicated the high thermostability of enzyme. We have modelled and performed dynamics simulation on apo and NADH bound *MtbDprE2* to understand the enzyme dynamics involved in NADH recognition. The NADH binding to *MtbDprE2* showed minor conformational changes in active site residues and orientation between and small and large domains in dynamics simulation. The knowledge of structure and dynamical aspect of NADH binding may contribute in specific inhibitors development of *Mtb*DrE2 enzyme.

## Abbreviations used

DprE2: decaprenylphosphoryl-β-D-ribose epimerase-2
DPR: decaprenylphosphoryl-β-D-ribofuranose
DPX: decaprenylphosphoryl-β-D-ribose
DPA: decaprenylphosphoryl-β-D-arabinofuranose
BTZ043: benzothiazinone
DNB: dinitrobenzamide
*E. coli*: *Escherichia coli*
LB: luria Bertani
PME: Partial Mesh Ewald
CD: Circular dichroism
PDB: Protein data bank

## Conflict of interest statement

None

## Acknowledgements

The partial funding from DST–PURSE, UPE-II, UGC resource networking and Department of Biotechnology (No. PAC-SLS-AKS-DBT-01131216-737) are gratefully acknowledged. JNU, AIRF facility for their kind support in conducting the CD experiment. ShantiPal and Arkita were supported by fellowship from UGC.

## Supplementary figures

**Figure S1.**
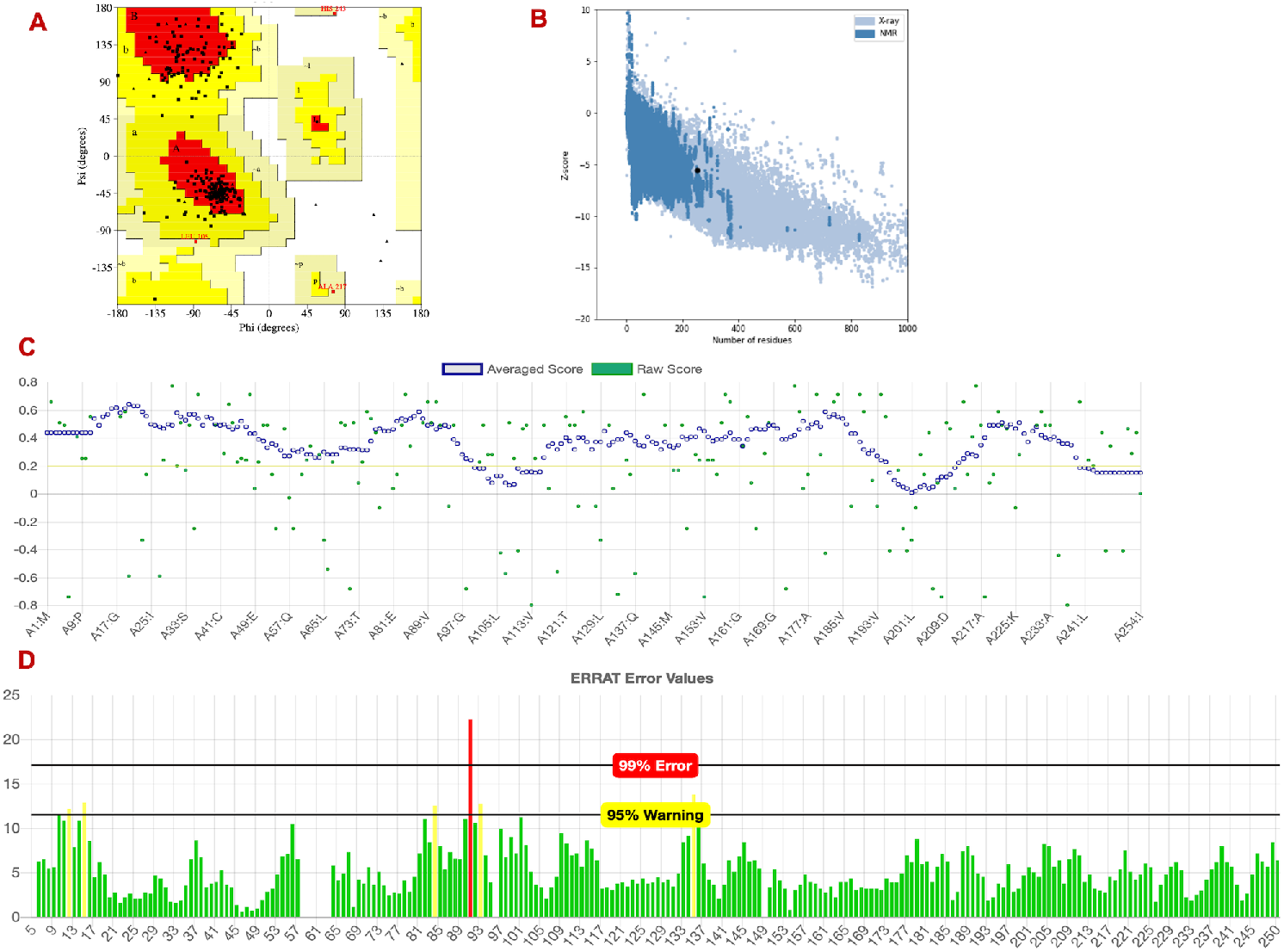
Quality check on apo-*MtbDprE2* model obtained after 1 ns of dynamics simulation. **(A)** The 95% residues in most favored and 5% residues in additionally allowed regions were obtained in the Ramachandran plot. **(B)** Z score of −5.5 was observed in ProSA plot, which indicate that overall quality of model is good. **(C)** the regions (yellow color) in model, that can be rejected with 95% confidence is shown in ERRAT plot. **(D)** the regions, whose threshold scores greater than 0.2 is shown in Verify 3D plot.

**Figure S2.**
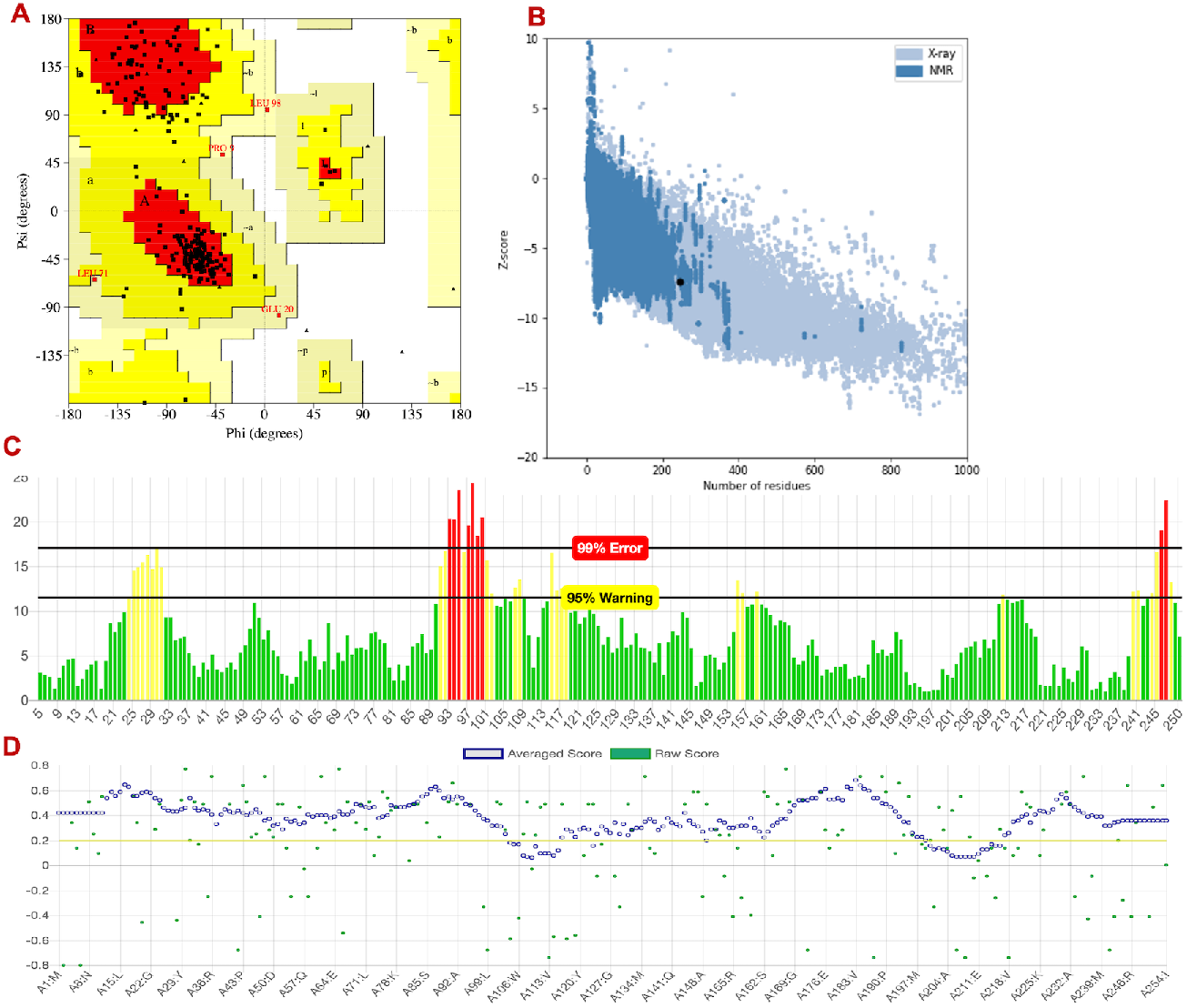
Quality check on *MtbDprE2*+NADH complex obtained after 1 ns of dynamics simulation. **(A)** 96% residues in most favored and 4% residues in additional allowed regions were observed in Ramachandran plot (**B**) Z score of – 7.36 was observed in ProSA plot, which indicate that model has good quality. **(C)** ERRAT plot shows the regions, that can be rejected with 95% confidence (yellow). **(D)** the regions whose threshold scores greater than 0.2 is shown in Verify 3D plot.

## Notes

### Competing Interest Statement

The authors have declared no competing interest.

